# Vocal fold injury alters acoustic phenotype despite preserved vocal activity

**DOI:** 10.64898/2026.09.29.755416

**Authors:** Shannon Hogan, Jung Keun Lee, Tobias Riede

## Abstract

Vocal communication depends on the biomechanical integrity of the vocal folds, yet how impairment alters vocal behavior remains poorly understood. We used the California mouse (*Peromyscus californicus*), a highly vocal species that produces songs through airflow-induced vocal fold vibration, to experimentally examine how defined vocal fold injury alters vocal production. Longitudinal recordings were combined with histological assessment to track changes in vocal function after impairment. Injury produced acute edema of the lamina propria and thyroarytenoid muscle. Despite stable levels of vocal activity, the vocal phenotype shifted characteristically following injury. Thus, vocal behavior can persist despite substantial alteration of the biomechanical tissues underlying sound production, demonstrating that maintenance of behavioral output does not necessarily indicate preservation of the physiological system that generates it. More broadly, these findings establish an experimental link between vocal fold injury and vocal phenotype and provide a framework for understanding how tissue damage and remodeling can reshape acoustic communication even when vocal behavior is maintained. Because repeated vocalization can impose substantial mechanical demands on vocal folds, this framework may have broad relevance for understanding how vocal use, injury, and tissue remodeling interact to shape vocal communication.

## Introduction

Animal behavior depends on the structural and physiological integrity of the systems that generate it. When these systems are impaired, behavior may change, yet it remains unclear whether a behavior can persist despite disruption of the physiological characteristics of the structures that produces it. Vocal communication provides a tractable example of this broader problem: vocalizations in many vertebrates are generated by airflow-induced vocal fold vibrations, and structural and mechanical properties of vocal folds shape the resulting acoustic signals (1–4). Vocal folds are exposed to repeated mechanical forces during vocalization especially in contexts involving frequent or intense vocal use, and can undergo injury and remodeling. Determining whether and how impairment of the vocal apparatus alters vocal behavior can reveal how changes to the physiological substrate of behavior become expressed as changes in its observable phenotype.

Understanding this relationship requires experimental systems in which tissue injury can be defined and behavioral consequences quantified. The California mouse (*Peromyscus californicus*) provides an opportunity to investigate this structure–function relationship because it produces highly stereotyped vocalizations (5,6). Long-distance vocalizations or songs are generated by airflow-induced vibration of the vocal folds, while coordinated movements of the vocal tract shape the acoustic signal (7–8). Although the California mouse larynx is broadly similar to that of the house mouse (*Mus musculus*), its vocal folds exhibit species-specific structural and cellular specializations (9–10). Its mode of vocal production, however, relies on airflow-induced vocal fold vibration, a mechanism broadly conserved among mammals, making it a useful model for investigating how tissue properties shape vocal function.

Here, we experimentally injured one vocal fold in California mice and examined vocal behavior following injury. We quantified vocal activity and acoustic song parameters and related these measures to histopathological changes in the vocal folds. We tested whether localized tissue injury alters vocal behavior and whether variation in vocal characteristics is associated with the extent of tissue damage or overall vocal activity. By experimentally linking tissue injury, vocal behavior, and subsequent tissue recovery, we investigate how structural changes in a biomechanical organ are reflected in animal behavior.

## Methods

### Animals

Captive-born male and female Peromyscus californicus were maintained at the Midwestern University animal facility. The breeding colony was established in 2019 from 12 founders obtained from the Peromyscus Genetic Stock Center (Columbia, SC, USA), whose stock derives from wild-caught animals. Mice were housed in polycarbonate cages (48 × 26.5 × 20 cm) with wood-chip bedding, paper nesting material, and plastic tubes under a 12 h:12 h light cycle (lights on at 1400 h), 21–23 °C, and 25–35% relative humidity, with standard rodent chow (Teklad 2918; Envigo, Madison, WI, USA) and water ad libitum. All procedures were approved by the Midwestern University Institutional Animal Care and Use Committee (Protocol 4112).

Sixty-eight mice (9–18 months) were included. Forty-seven (24 males, 23 females) received unilateral right vocal fold injury and six (3 males, 3 females) sham surgery. Of the injured mice, 13 (7 males, 6 females) were euthanized within 24 h for acute histopathological analysis, while 34 recovered for up to 4 weeks as the chronic injury group. Vocal behavior was monitored longitudinally in chronic and sham groups, and chronic vocal phenotype was related to vocal fold structure assessed at the end of recording. Thirty-two chronic-injury mice provided sufficient longitudinal acoustic data for recovery analyses. Histological comparisons additionally included 15 uninjured mice (7 males, 8 females) from previous studies (10).

### Surgical procedure of the vocal fold injury

Unilateral vocal fold injury was induced by mild mechanical compression of the right vocal fold under ketamine–xylazine anesthesia (80 and 8 mg kg ¹, respectively). Anesthetic depth was confirmed by respiratory rate, absence of the pedal withdrawal reflex, and reduced whisker movement; no supplemental anesthesia was required. The larynx was exposed and the glottis visualized under an operating microscope. The right vocal fold was compressed for ∼2 s using a 23-gauge intraocular micro-alligator clamp, while the left remained unmanipulated as an internal control (Fig. 1a). Sham-operated animals underwent identical anesthesia and glottic exposure without vocal fold manipulation. Postoperatively, animals received a single subcutaneous dose of Ringer’s lactate and carprofen (Rimadyl)/enrofloxacin (Baytril) tablets (on postoperative days1 – 3) (Bio-Serv, MD150-050) as routine care.

**Figure 1.**
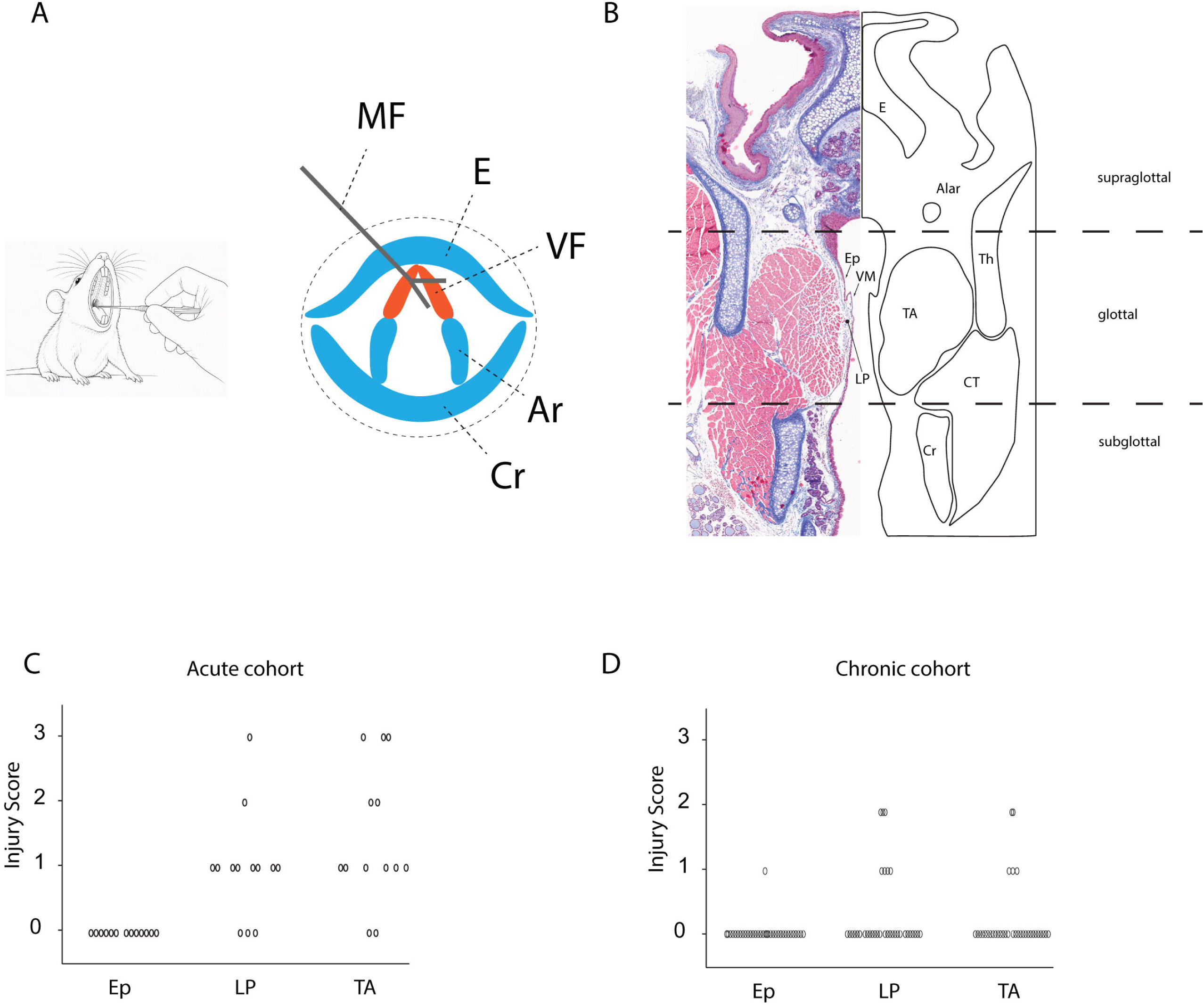
Schematic representation and histological anatomy of the California mouse larynx, and severity of vocal fold injury following surgery. **(A)** Vocal folds were visualized perorally. Schematic image of the intra-oral view of the larynx showing the site of injury, which was created by compressing the right vocal fold with microforceps (MF) under direct visualization. **(B)** Coronal schematic illustrating the major laryngeal structures and midmembranous section of the larynx stained with Masson’s trichrome. **(C–D)** Distribution of vocal fold injury scores in acute **(C**; *n* = 13; assessed within 24 h after surgery) and chronic **(D**; *n* = 32; assessed 14–30 days after surgery) samples. Injury severity was scored on a scale of 0–3 according to the criteria described in Table 1 and exemplified in figure 2. Each point represents an individual animal. No epithelial injury was observed in either group. Injury was primarily localized to the lamina propria and thyroarytenoid (TA) muscle and was more severe in the acute than in the chronic group. Abbreviations: MF, microforceps; E, epiglottis; VF, vocal fold; Ar, arytenoid cartilage; Cr, cricoid cartilage; Th, thyroid cartilage; TA, thyroarytenoid muscle; CT, cricothyroid muscle; Ep, vocal fold epithelium; LP, lamina propria; VM, vocal membrane. Mouse image in panel A was generated using Microsoft Copilot (AI-generated) and modified by the authors.

**Table 1:** Histological scoring system for evaluation of vocal fold injury after unilateral mechanical damage. A semi-quantitative scoring system was established to characterize injury severity and tissue remodeling in the epithelium, lamina propria, and thyroarytenoid muscle. Each anatomical layer was scored independently from 0 (normal) to 3 (severe injury) based on the extent of epithelial damage, lamina propria expansion, inflammatory infiltration, fibrosis, myofiber degeneration, necrosis, muscle atrophy, and hemorrhage. Lamina propria thickness was assessed relative to normal reference measurements. Acute injury-associated features were evaluated at 24 h post-injury, whereas chronic remodeling features, including fibrosis and muscle atrophy, were evaluated at 4 weeks post-injury.

| Score | Epithelium | Lamina propria | Muscle |
| --- | --- | --- | --- |
| Normal (0) | Within normal limits | Within normal limits | Within normal limits |
| Mild (1) | Focal epithelial degeneration or disruption involving fewer than 5 epithelial cells, without ulceration or loss of epithelial continuity. | Mild expansion due to edema, with occasional leukocyte infiltration. Fibrous tissue deposition may be present in chronic samples. | Mild edema within the perimysium. Myofiber changes, including swelling, vacuolization, and hypereosinophilia, involve up to 10% of myofibers. Mild muscle atrophy may be present in chronic samples. |
| Moderate (2) | Multifocal epithelial degeneration or disruption involving more than 5 epithelial cells, with maintained epithelial continuity and no extensive ulceration. | Expanded by edema with up to 50% increase in thickness compared with normal reference values. Moderate leukocyte infiltration is present. Fibrous tissue deposition is present in chronic samples. | Moderate edema within the perimysium with myofiber degeneration, including swelling, vacuolization, and hypereosinophilia involving more than 10% and up to 50% of myofibers. Muscle atrophy is present in chronic samples. |
| Severe (3) | Extensive multifocal epithelial loss with ulceration or marked disruption of epithelial continuity involving large areas of the epithelial surface. | Markedly expanded by edema with more than 50% increase in thickness compared with normal reference values. Dense leukocyte infiltration is present. Extensive fibrous tissue deposition is present in chronic samples. | Severe edema within the perimysium with extensive myofiber degeneration, including swelling, vacuolization, and hypereosinophilia involving more than 50% of myofibers. Necrotic myofibers may be present, characterized by fragmentation, hypereosinophilic cytoplasm, and centralized nuclei. Leukocyte infiltration and/or hemorrhage may be present in acute samples. Severe muscle atrophy may be present in chronic samples. |

### Histology acquisition, injury scoring, and morphometric analysis

Following euthanasia, larynges were fixed in 10% neutral buffered formalin, paraffin-embedded, serially sectioned transversely (5 µm) throughout the glottal region at 50–100 µm intervals, and stained with hematoxylin and eosin or Masson’s trichrome. Images were acquired using ImageScope (Leica Biosystems).

A blinded, board-certified pathologist evaluated injury and developed a study-specific scoring system based on histopathological features in all 68 mice and published criteria (Table 1). Mid-membranous glottal sections were scored separately for epithelium, lamina propria, and thyroarytenoid (TA) muscle (Fig. 1b), each from 0 (normal) to 3 (severe), for control (n = 15), sham (n = 6), acute-injury (n = 13), and chronic-injury (n = 34) groups. Layer scores were summed to yield a composite score (maximum = 9).

Because the TA muscle showed the primary structural alterations, morphometry compared injured right and contralateral left TA muscles. Cranial–caudal (CC) distance, medial–lateral (ML) distance, muscle cross-sectional area, and mean myofiber cross-sectional area were quantified. CC distance spanned the most caudal to cranial muscle boundaries; ML distance was measured perpendicular to the CC axis at maximal muscle thickness; muscle area was manually segmented; and myofiber area was calculated from individual fibers using ImageScope annotation tools.

To establish myofiber sampling adequacy, 200 fibers from four vocal folds across four randomly selected specimens were analyzed; cumulative mean analysis indicated stabilization after ∼50 fibers. To minimize size-based sampling bias, TA cross-sections were divided into eight similar-sized transects, and the four largest and four smallest fibers in each transect were measured (64 fibers/vocal fold). This yielded the lowest intra-observer variability (CV ≈1% across four repeated measurements in one vocal fold) and was used for all subsequent analyses.

### Triggering singing and vocal monitoring before and after injury

Preliminary work showed that spontaneous singing in solitary-housed animals can be triggered by auditory contact with established pair-mates. Chronic-injury and sham-operated animals were established male–female pairs that had previously produced ≥1 litter. To induce separation-associated singing, pairs were housed individually in transparent polycarbonate cages positioned side-by-side, allowing auditory and limited visual but not physical contact; pair members remained within acoustic range throughout recording under identical sensory conditions. Female reproductive status was not determined. Animals had no prior recording or experimental exposure. Recordings were conducted in the standard housing room under baseline husbandry conditions, and all animals were euthanized at study completion.

Chronic-injury (N = 34) and sham-operated (N = 6) mice were monitored continuously (24 h/day) for 2 weeks before surgery to establish baseline vocal behavior, account for daily variability, and estimate normal acoustic ranges, then for up to 4 weeks postoperatively. Vocalizations were recorded with an ultrasonic microphone (CM16/CMPA, Avisoft Bioacoustics, Glienicke, Germany) ∼5 cm above each cage, digitized at 300 kHz using an UltraSoundGate 416Hb (Avisoft Bioacoustics), and stored uncompressed with Avisoft Recorder. Recordings included spontaneous vocalizations and cage noise; all files were visually confirmed for songs.

Song recordings included 40 mice (34 injured, 6 sham), yielding 14,474 calls and 40,248 syllables. Sham mice contributed 2,594 calls and 7,244 syllables before and after surgery. Injured mice contributed 11,880 calls and 33,004 syllables before and after injury, respectively; 32 of 34 injured mice contributed post-injury recordings.

### Characterization of vocal activity and acoustic parameters

Because injury-induced sickness behavior can reduce activity and social interaction and thereby confound acoustic measures of vocal-fold function (11, 12), vocal activity was quantified as the number of songs produced per 24-h recording period (one complete dark–light cycle). To distinguish overall activity from vocal-bout structure, syllables per song were also quantified; California mouse songs comprise 1–8 syllables (5, 6).

Seven acoustic parameters were measured for each syllable. Fundamental frequency (f□) and sound intensity were extracted at 10-ms intervals using a custom Praat script and used to calculate mean, minimum, and maximum f□, syllable duration, syllable repetition rate, mean sound intensity, and centroid dispersion (CD).

CD quantified acoustic variation within two-dimensional [f□, I] space. For each recording period (pre/post injury or sham operation), the centroid was calculated from all tracked [f□, I] points, and CD was defined as the mean Euclidean distance of tracking points from that centroid:

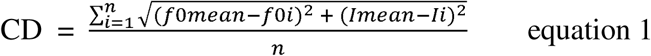

where (*f* _mean_) and (I_mean_) are the period-wide mean *f* and intensity, *f _i_* and *I_i_* are instantaneous values at tracking point i, and n is the total number of tracking points. Higher CD indicates greater dispersion and a broader acoustic range; lower CD indicates more restricted vocal output.

### Data analysis

#### a) Characterization of the vocal fold injury

Histological injury scores for the epithelium, lamina propria, and muscle were treated as ordinal data. Group differences among control, acute-injury, chronic-injury, and sham-operated animals were assessed using nonparametric tests, with post hoc pairwise comparisons by Wilcoxon rank-sum tests; effect sizes were quantified using rank-biserial correlation. Associations between lamina propria and thyroarytenoid muscle injury scores were assessed using Kendall’s and Spearman’s rank correlations separately for acute, chronic, and control groups. One-sided tests were used for the hypothesized positive associations between tissue compartments.

Right–left asymmetry in thyroarytenoid morphology was assessed within subjects. For each morphometric variable (cranial–caudal length, medial–lateral width, thyroarytenoid muscle cross-sectional area, myofiber cross-sectional area, and CC:ML ratio), right–left differences were calculated per animal. Normality of paired differences was assessed using the Shapiro–Wilk test, followed by paired t-tests or Wilcoxon signed-rank tests as appropriate. Both tests were computed for each comparison, with inference based on distributional assumptions.

#### b) Phenotyping singing behavior after vocal fold injury

To test whether injury altered vocal behavior, vocal activity and syllables/song were compared within animals before and after injury or sham surgery using paired tests. Acoustic changes were assessed at three levels: (1) individual acoustic parameters, (2) multivariate vocal phenotype, and (3) nonlinear vocal phenomena (NLPs), reflecting sound-source instability.

Seven acoustic parameters were compared within animals between pre- and postoperative recordings, with false discovery rate (FDR) correction where appropriate. For multivariate analysis, the seven parameters were standardized across syllables and subjected to principal component analysis (PCA). PC scores were averaged across syllables within each mouse and condition to generate mouse-level phenotypes. PC1 and PC2, representing the major axes of acoustic variation, were used for visualization, with loadings reported to characterize parameter contributions. For pre-injury, post-injury, pre-sham, and post-sham conditions, mouse-level PC1/PC2 centroids were calculated. Euclidean distances between pre- and postoperative centroids quantified group-level phenotype shifts, while mean distances of mice from their centroids quantified within-group dispersion. Paired changes in PC scores assessed within-animal shifts, and Hotelling’s T² tested multivariate effects of injury; sham animals were analyzed in parallel to identify surgery-related changes.

For visualization, each animal’s pre/post PC1/PC2 coordinates were connected by a trajectory and translated so its pre-treatment coordinate aligned with the corresponding condition centroid. The same translation vector was applied to its post-treatment coordinate, preserving trajectory length, angle, and direction while changing only absolute position.

NLPs (subharmonics, biphonation, chaos, and frequency jumps) were quantified as the proportion of syllables exhibiting each feature. Pre/post changes were assessed within individuals using paired statistics, with sham animals analyzed in parallel to distinguish injury effects from procedural artifacts.

#### c) Characterization of vocal recovery and its association with histological injury score and post-operative vocal activity

Vocal recovery was quantified as the distance of each postoperative syllable from its corresponding baseline position in PCA space, with lower values indicating greater similarity to the pre-injury phenotype. Temporal recovery was tested using a linear mixed-effects model with postoperative day as a fixed effect and mouse as a random intercept and slope, allowing individuals to differ in baseline distance and recovery rate. Models were fitted using REML with Satterthwaite-approximated degrees of freedom. The postoperative-day coefficient represented the population-average daily change in acoustic distance; negative values indicated progressive recovery. Ninety-five percent confidence intervals (CIs) were calculated using Wald intervals. Population-level predictions and 95% confidence bands were generated from fixed effects and plotted with individual observations and mouse-specific trajectories.

The population postoperative-day coefficient was interpreted as the average recovery rate (change in acoustic distance/day), rather than as a separate hypothesis test. Mouse-specific slopes were calculated by adding each estimated random-slope deviation to the fixed postoperative-day coefficient; negative, near-zero, and positive slopes indicated decreasing, stable, and increasing acoustic distance, respectively. These slopes were summarized descriptively and compared with histological scores.

Associations between end-point histological injury score and recovery were evaluated in the vocal-activity models described below. The score main effect tested whether injury score predicted overall acoustic distance after accounting for postoperative day and vocal activity, whereas the postoperative day × score interaction tested whether score predicted recovery rate. Thus, the main effect represented differences in distance at the reference postoperative day and centered covariates, while the interaction represented differences in temporal recovery rate.

Postoperative vocal activity was quantified for each injured mouse from day 1 through the final recording day. A complete calendar-day sequence was generated for each mouse, daily vocal counts were joined to it, and days without vocalizations were assigned zero. Mean activity across 32 mice was strongly right-skewed and was therefore natural-log transformed and standardized to a z-score. The activity coefficient thus represented the expected change in acoustic distance per 1-SD increase in log-transformed activity.

Vocal activity was tested for associations with recovery using two models: the activity main effect tested whether activity predicted overall acoustic distance after accounting for postoperative day and histological score, and the postoperative day × activity interaction tested whether activity predicted recovery rate. Both models included mouse random intercepts and slopes. The primary activity test was the day × activity interaction: negative values indicated faster recovery among more active mice and positive values slower recovery. The day × histological score interaction was included to assess whether injury score modified recovery rate. Both models used REML, Satterthwaite degrees of freedom, and 95% Wald CIs for primary activity effects.

Analyses were conducted in R 4.5.2. Tests were two-tailed unless otherwise specified, with α = 0.05; multiple-comparison correction was applied within each test family where applicable.

## Results

### Characterization of the vocal fold injury

Acute injury primarily involved the lamina propria and thyroarytenoid (TA) muscle, with preserved epithelial integrity (Fig. 2). Lamina propria lesions included edema and inflammatory cell infiltration, whereas TA muscle injury was characterized by myofiber swelling, vacuolization, hypereosinophilia, and severe myofiber degeneration (Fig. 2). Representative moderate-to-severe injury scores are shown in Supplemental Fig. 1a–c.

**Figure 2.**
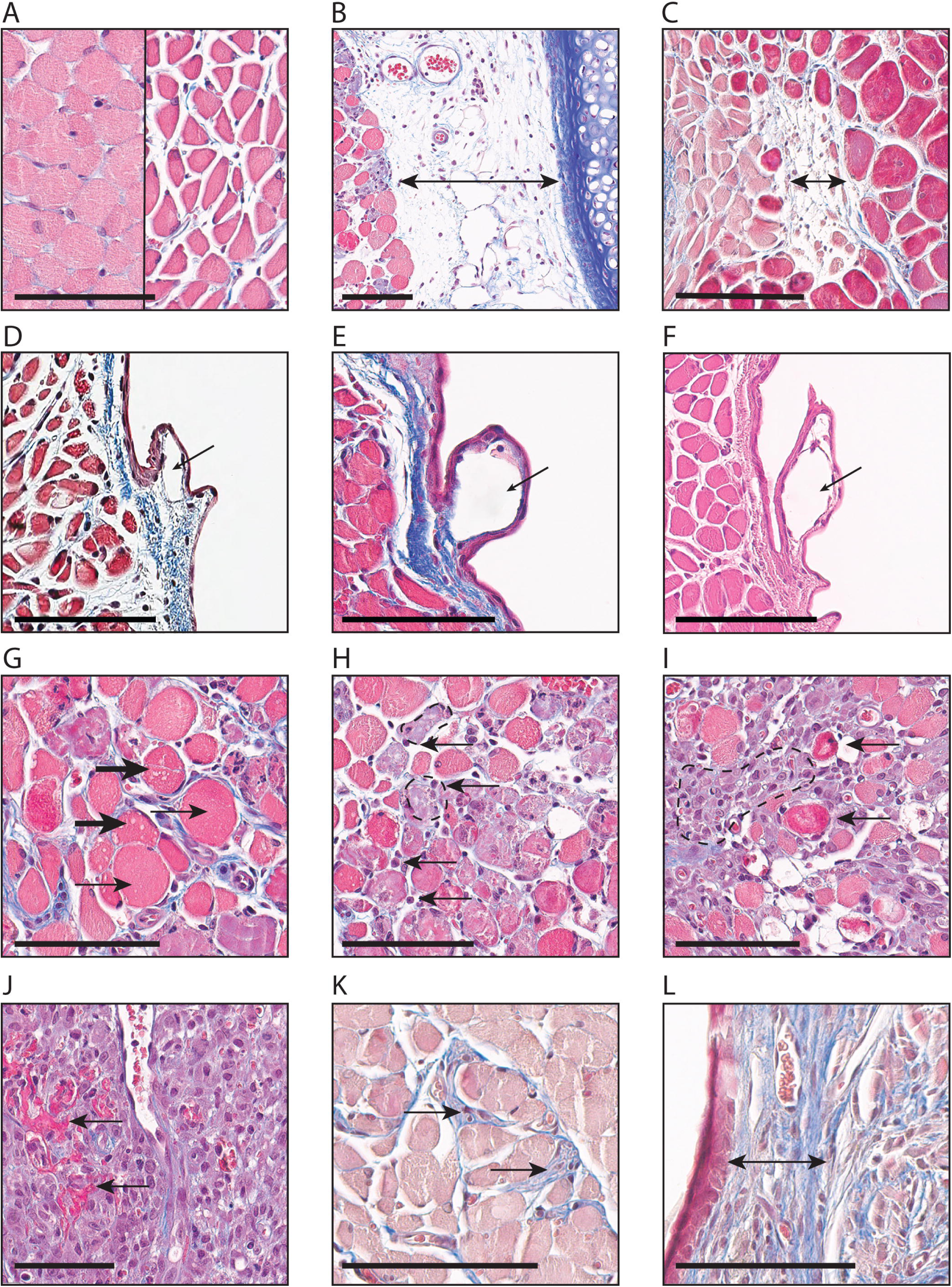
Histological signs of pathology in the vocal folds of California mice. Coronal sections of the larynx stained with hematoxylin and eosin (H&E) or Masson’s trichrome are shown. (**A**–**C**) Representative examples of edema located between myofibers. **A**: Comparison of normal, densely packed myofibers (left) and widened edematous spaces between myofibers (right). **B**: Arrow indicates edema between the thyroarytenoid (TA) muscle and the thyroid cartilage. **C**: Edema within the TA muscle. (**D**–**F**) The vocal membrane is a small protrusion on the medial surface of each vocal fold composed of two layers of epithelium, a small amount of extracellular matrix, and a lymphatic vessel (arrow) (**D**). Lymphatic vessels within the vocal membrane exhibited varying degrees of dilation (arrows) in acute (**E**) and chronic (**F**) samples. (**G**–**L**) Representative myofiber pathologies included vacuolization (thick arrow) and severely swollen degenerated cytoplasm (thin arrow) (**G**); a degenerating necrotic myocytes (dashed outline) with infiltrating inflammatory cells (arrows) (**H**); a completely degenerated necrotic myocyte with hypereosinophilia (arrows, **I**), hemorrhage (arrows, **J**); fibrosis between myofibers (**K**); and fibrosis within the lamina propria (**L**). Scale bars = 100 μm in all panels.

Injury scores were substantially higher in the acute group than in the chronic or control groups (Fig. 1c,d; Table 2). The greatest differences were observed for lamina propria and TA muscle injury, whereas epithelial injury was minimal across all groups (Table 2). Direct comparison of acute and chronic groups demonstrated significantly higher lamina propria scores in the acute group (Wilcoxon rank-sum test, *W* = 339, *p* = 0.0010). The corresponding rank-biserial correlation was 0.48 (95% CI, 0.20–0.72). Muscle injury scores were also significantly higher in acute than the chronic group (*W* = 380, *p* < 0.001), with a large rank-biserial correlation (0.66, 95% CI, 0.43–0.86).

**Table 2.** Injury scores by experimental group. Injury scores are presented as mean ± standard deviation (SD). AC, acute injury group (1 day post-injury); CH, chronic group (2 to 4 weeks after injury); CO, control group. E, epithelium; LP, lamina propria; M, thyroarytenoid (TA) muscle. *n* indicates the number of animals in each group.

| Group | n | E | LP | M |
| --- | --- | --- | --- | --- |
| AC | 13 | 0 ± 0 | 1.00 ± 0.82 | 1.46 ± 1.05 |
| CH | 34 | 0.029 ± 0.171 | 0.294 ± 0.629 | 0.206 ± 0.538 |
| CO | 16 | 0 ± 0 | 0.188 ± 0.403 | 0.125 ± 0.342 |

Lamina propria and muscle injury scores were strongly positively associated in the acute group. Kendall’s rank correlation yielded τ = 0.747 (*p* = 0.0017, one-sided), while Spearman’s rank correlation yielded ρ = 0.795 (*p* = 0.0006, one-sided). Thus, greater lamina propria injury was associated with greater TA muscle injury in the acute group. This association was weak in the chronic group, with no statistically significant correlation (Kendall’s τ = 0.207, *p* = 0.108; Spearman’s ρ = 0.213, *p* = 0.114). Similarly, no positive association was detected in control animals (Kendall’s τ = −0.182, *p* = 0.759; Spearman’s ρ = −0.182, *p* = 0.750).

Morphometric analysis revealed a significantly larger TA muscle cross-sectional area on the injured side in the acute group (*p* = 0.036) and significantly shorter CC-distance on the injured side in the chronic group (*p* = 0.007; Table 3). No other significant right–left differences were observed in either group. Control animals showed no significant right–left differences for any morphometric parameter (Table 3).

**Table 3.** Right–left morphometric comparisons by experimental group. Values are presented as mean ± SD. Right VF denotes the injured side and left VF the contralateral side. AC, acute injury group; CH, healing group; CO, control group; TA, thyroarytenoid; CC, cranial-caudal; ML, medial-lateral. *P* values are from paired *t*-tests; *p* < 0.05 indicates a significant right–left difference.

| Group | Morphometric parameter | Right VF | Left VF | P value |
| --- | --- | --- | --- | --- |
| **AC** | TA muscle cross-sectional area ($\mu\text{m}^2$) | $455 \pm 120 \times 10^3$ | $408 \pm 83 \times 10^3$ | **0.036** |
| | Myofiber cross-sectional area ($\mu\text{m}^2$) | $392.6 \pm 92.6$ | $364.8 \pm 41.8$ | 0.223 |
| | CC length ($\mu\text{m}$) | $900.2 \pm 148.5$ | $906.9 \pm 125.6$ | 0.803 |
| | ML width ($\mu\text{m}$) | $647.3 \pm 114.6$ | $626.0 \pm 114.6$ | 0.569 |
| **CH** | TA muscle cross-sectional area ($\mu\text{m}^2$) | $451,647 \pm 103,538$ | $448,586 \pm 79,028$ | 0.800 |
| | Myofiber cross-sectional area ($\mu\text{m}^2$) | $334.5 \pm 65.2$ | $345.4 \pm 50.0$ | 0.697 |
| | CC length ($\mu\text{m}$) | $920.6 \pm 140.2$ | $975.0 \pm 96.8$ | **0.007** |
| | ML width ($\mu\text{m}$) | $615.4 \pm 103.0$ | $590.6 \pm 69.6$ | 0.123 |
| **CO** | TA muscle cross-sectional area ($\mu\text{m}^2$) | $369,226 \pm 46,724$ | $364,281 \pm 46,873$ | 0.270 |
| | Myofiber cross-sectional area ($\mu\text{m}^2$) | $366.9 \pm 60.0$ | $353.0 \pm 51.3$ | 0.151 |
| | CC length ($\mu\text{m}$) | $915.4 \pm 73.6$ | $910.3 \pm 71.8$ | 0.820 |
| | ML width ($\mu\text{m}$) | $533.2 \pm 42.3$ | $526.5 \pm 59.2$ | 0.320 |
| | CC:ML ratio | $1.709 \pm 0.203$ | $1.743 \pm 0.243$ | 0.339 |

Together, these findings demonstrate acute injury was confined primarily to the lamina propria and TA muscle, with epithelial integrity preserved. Acute histopathologic injury resolved after 2 to 4 weeks. Reduced lamina propria and muscle injury scores, together with loss of their strong acute association, indicate resolution of the coordinated injury response following healing. Persistent reduction in TA muscle CC distance despite preserved cross-sectional area suggests residual structural remodeling. Controls showed no systematic right–left morphometric differences.

### Phenotyping singing behavior after vocal fold injury Vocal activity

Daily vocal activity varied substantially among animals. In sham-operated controls (*n* = 6), mean vocal activity decreased following surgery, from 72.9 ± 67.5 to 31.0 ± 21.8 songs/24 h, but this change was not statistically significant (paired *t*-test, *t*(5) = −1.45, *p* = 0.206).

In the chronic injury cohort (*n* = 34), mean vocal activity likewise decreased following unilateral vocal fold injury, from 16.2 ± 24.8 to 12.5 ± 12.3 songs/24 h. Two animals produced no vocalizations after injury. Overall, vocal activity did not change significantly following injury (paired *t*-test, *t*(31) = −0.94, *p* = 0.354; mean change = −4.73 calls/24 h, 95% CI, −15.20 to 5.74). Because individual songs may comprise one or multiple syllables, we also examined the number of syllables per song. Among animals with songs recorded before and after injury (*n* = 32), the mean number of syllables per song remained essentially unchanged following injury, from 2.61 ± 0.71 before injury to 2.63 ± 0.66 after injury (mean change = 0.019 syllables/song, 95% CI, −0.157 to 0.194). This difference was not statistically significant (paired *t*-test, *t*(31) = 0.219, *p* = 0.828). Similarly, sham-operated animals showed no significant change in the mean number of syllables per song, from 3.24 ± 0.89 before surgery to 2.98 ± 1.00 after surgery (mean change = −0.267 syllables/song, 95% CI, −1.285 to 0.752; paired *t*-test, *t*(5) = −0.673, *p* = 0.531). Thus, unilateral vocal fold injury did not produce a detectable change in either overall daily vocal activity or the number of syllables produced per song.

### Acoustic parameters

Vocal acoustic parameters were stable following sham surgery, with no significant pre–post changes detected for any of the seven measures after correction for multiple comparisons (Table 4). In contrast, vocal fold injury produced coordinated changes across multiple dimensions of vocal output. Mean, maximum, and minimum fundamental frequency, mean intensity, and syllable duration all decreased significantly following injury (all FDR-adjusted *P* < 0.001), whereas centroid dispersion and syllable repetition rate increased (FDR-adjusted *P* = 0.041 and 0.003, respectively; Table 4). Descriptive values (mean ± SD) for all acoustic parameters before and after sham surgery or injury are provided in Supplemental Table 2.

**Table 4:**
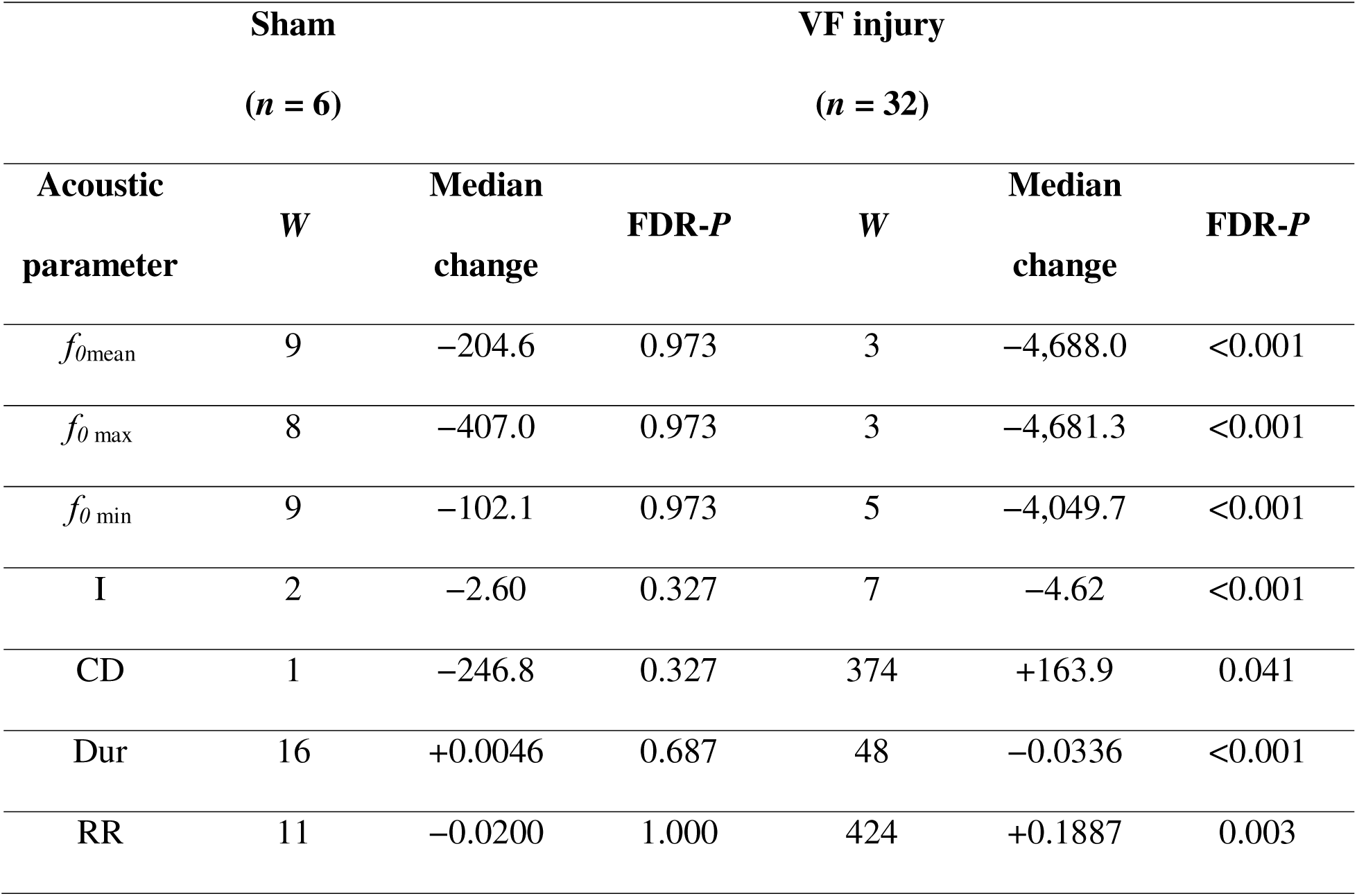
Pre–post changes in vocal acoustic parameters following sham surgery and chronic vocal fold injury. Values represent paired Wilcoxon signed-rank tests comparing recordings obtained before and after surgery or injury. Median change is calculated as post-intervention minus pre-intervention values; negative values indicate a decrease following intervention. *P* values were adjusted for multiple comparisons using the false discovery rate (FDR) procedure across the seven acoustic parameters within each group. *f_0_*, fundamental frequency; I, mean intensity; CD, centroid dispersion; Dur, syllable duration; RR, syllable repetition rate.

To determine whether these changes collectively represented a shift in the overall vocal phenotype, we used principal component analysis (PCA) to visualize the multidimensional acoustic space (Fig. 3 J-L). The first two principal components captured the major structure of variation, with PC1 dominated by fundamental-frequency measures and PC2 reflecting variation in intensity, centroid dispersion, duration, and repetition rate (Table 5). Pre- and post-sham recordings substantially overlapped in acoustic space, whereas post-injury recordings showed a pronounced displacement from the pre-injury phenotype (Fig. 3J). The displacement of the post-injury centroid from the pre-injury centroid was 2.38 PC units, compared with 0.26 PC units following sham surgery, representing an approximately 9-fold greater shift. The injury-associated displacement occurred predominantly along PC1 (ΔPC1 = −2.35), consistent with the prominent reduction in fundamental frequency. Within-group dispersion was similar across conditions (mean distance from centroid, 1.47–1.71 PC units), indicating that the injury-associated shift reflected a directional change in acoustic phenotype rather than increased heterogeneity. A multivariate test of within-animal changes confirmed a significant overall shift in acoustic phenotype following injury (Hotelling’s *T*² = 97.2, *P* < 0.001).

**Figure 3.**
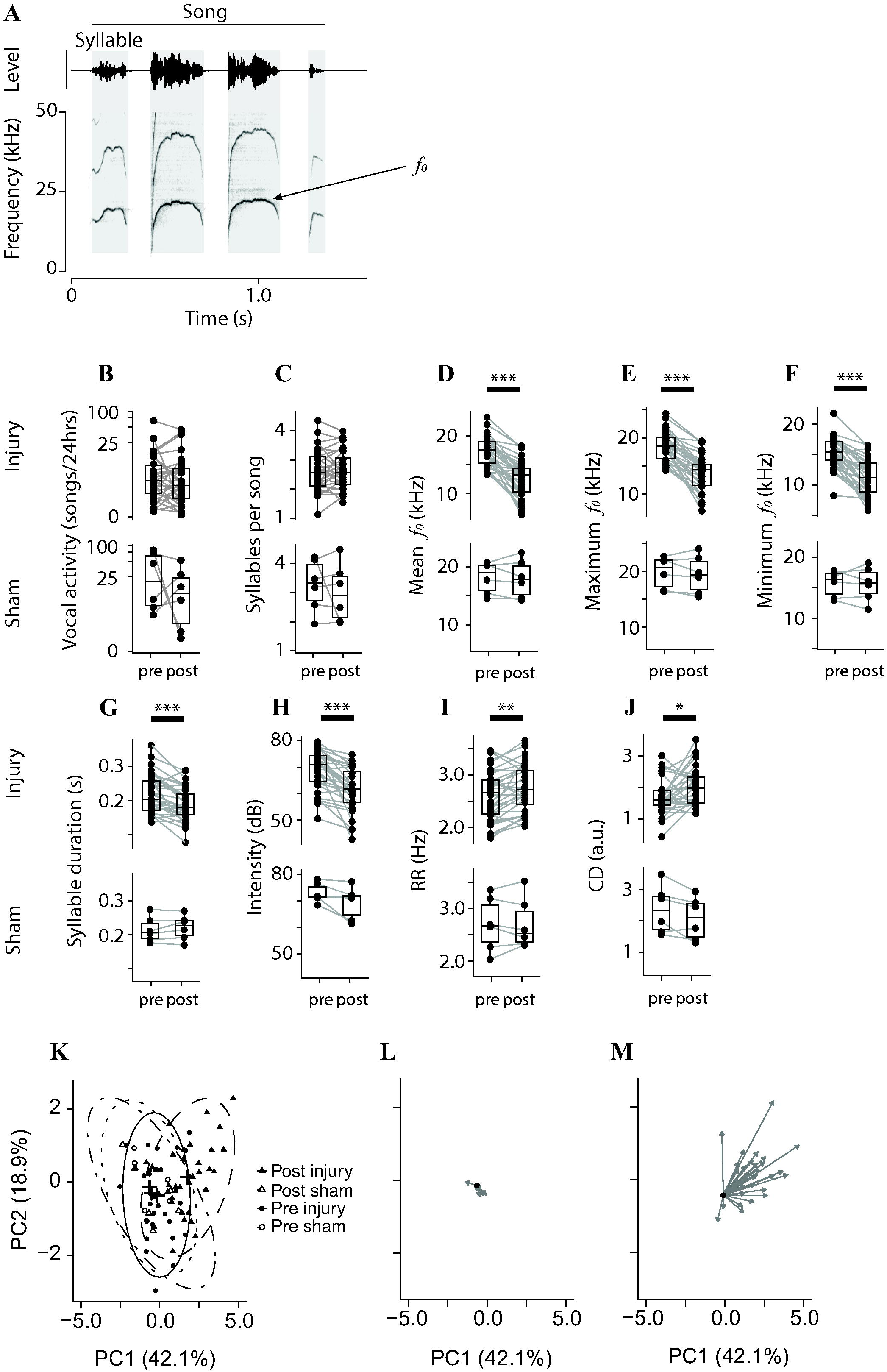
Vocal phenotype of California mouse songs before (“pre”) and after (“post”) unilateral vocal fold injury or sham surgery. (**A**–**J**) Vocal activity, syllables per call, and seven acoustic parameters were quantified in the chronic injury cohort (N = 32 mice) and sham-operated cohort (N = 6 mice). (**K**) Principal component analysis (PCA) of the seven acoustic variables (F0mean, F0min, F0max, mean intensity, duration, spectral slope, and repetition rate), with all pre- and post-treatment observations from injured and sham-operated animals included in a single standardized PCA. Each point represents one mouse; symbols indicate experimental condition and time point. Ellipses represent 95% confidence regions, and crosses indicate group centroids. The substantial overlap among pre- and post-sham observations and pre-injury observations indicates that anesthesia and the surgical procedure alone did not substantially alter song acoustics. (**L**) Individual sham-animal trajectories from pre- to post-surgery in PC1–PC2 acoustic space. (**M**) Individual injury-animal trajectories from pre- to post-injury. For visualization of individual trajectories, each animal’s pre-treatment position was translated to the centroid of its corresponding pre-treatment group, and the same translation was applied to its post-treatment position, preserving the original trajectory length, angle, and direction. Arrows indicate the direction of change from pre- to post-treatment. PC1 and PC2 axes are identical across panels **K**–**M**.

**Table 5:**
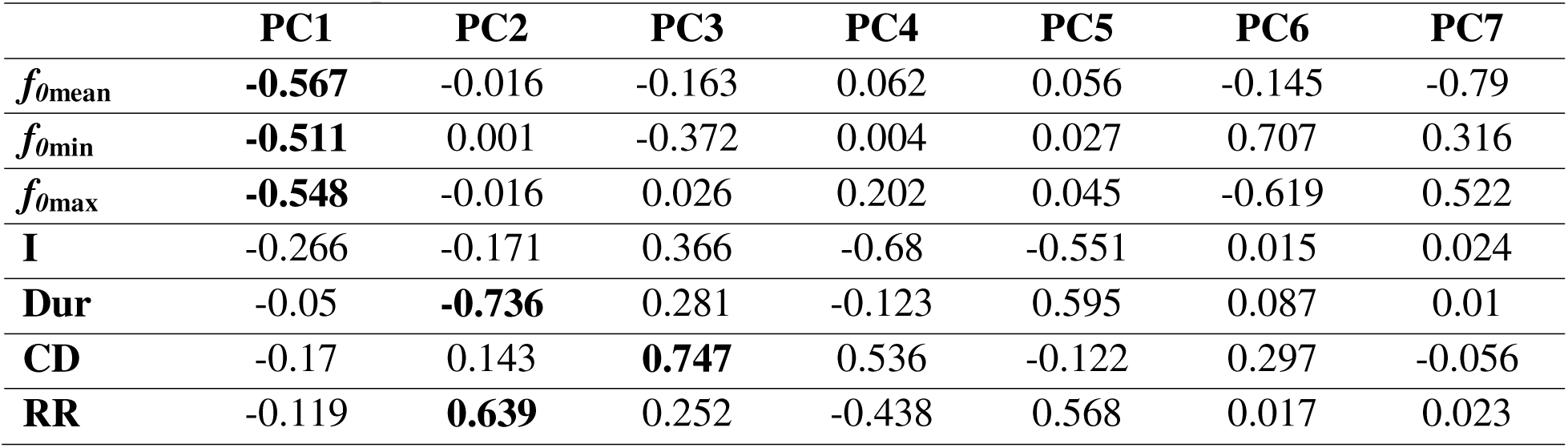
Principal component loadings for standardized acoustic variables. Loadings are shown for the seven acoustic variables included in the combined PCA of pre- and post-injury and sham observations. PC1–PC4 collectively explained ≥90% of total variance and were retained for subsequent analyses. Larger absolute values indicate stronger contributions of individual variables to each component.

|  | **PC1** | **PC2** | **PC3** | **PC4** | **PC5** | **PC6** | **PC7** |
| --- | --- | --- | --- | --- | --- | --- | --- |
| $f_{0\text{mean}}$ | **-0.567** | -0.016 | -0.163 | 0.062 | 0.056 | -0.145 | -0.79 |
| $f_{0\text{min}}$ | **-0.511** | 0.001 | -0.372 | 0.004 | 0.027 | 0.707 | 0.316 |
| $f_{0\text{max}}$ | **-0.548** | -0.016 | 0.026 | 0.202 | 0.045 | -0.619 | 0.522 |
| **I** | -0.266 | -0.171 | 0.366 | -0.68 | -0.551 | 0.015 | 0.024 |
| **Dur** | -0.05 | **-0.736** | 0.281 | -0.123 | 0.595 | 0.087 | 0.01 |
| **CD** | -0.17 | 0.143 | **0.747** | 0.536 | -0.122 | 0.297 | -0.056 |
| **RR** | -0.119 | **0.639** | 0.252 | -0.438 | 0.568 | 0.017 | 0.023 |

Individual trajectories in multivariate acoustic space revealed the direction and magnitude of within-animal changes from pre-to post-treatment (Fig. 3K, L). In sham animals (Fig. 3K), trajectories were generally short and variably directed indicating no consistent shift in acoustic phenotype after surgery. In contrast, injured animals exhibited larger and more consistent displacement in acoustic space following injury (Fig. 3L), Trajectories predominantly oriented toward the positive ends of PC1 and PC2. Thus, vocal fold injury produced coordinated changes across individual acoustic features that collectively displaced the vocal phenotype in multidimensional acoustic space, whereas sham surgery produced minimal displacement without increased within-group variability.

### Nonlinear phenomena (NLP)

The proportion of syllables exhibiting NLPs increased significantly following vocal-fold injury (n = 32 paired mice; Wilcoxon signed-rank test, V = 385, p = 0.0075; Fig. 4a). The median paired change in NLP rate was +0.154, corresponding to an approximately 15 percentage-point increase in NLP-containing syllables following injury. In contrast, sham-operated animals (n = 6 paired mice) did not show a significant change in NLP rate (V = 5, p = 0.295). The magnitude of the injury-associated increase varied considerably among individual animals, with a median change of 0.154 and a range of −0.618 to +0.872. Most interesting was that the rate of NLP syllables with subharmonics decreased, whereas biphonation and chaos increased. Frequency jumps showed relatively little change (Fig. 4a). Supplemental Fig. 2a–c shows songs from three example mice with various types of NLPs.

**Figure 4.**
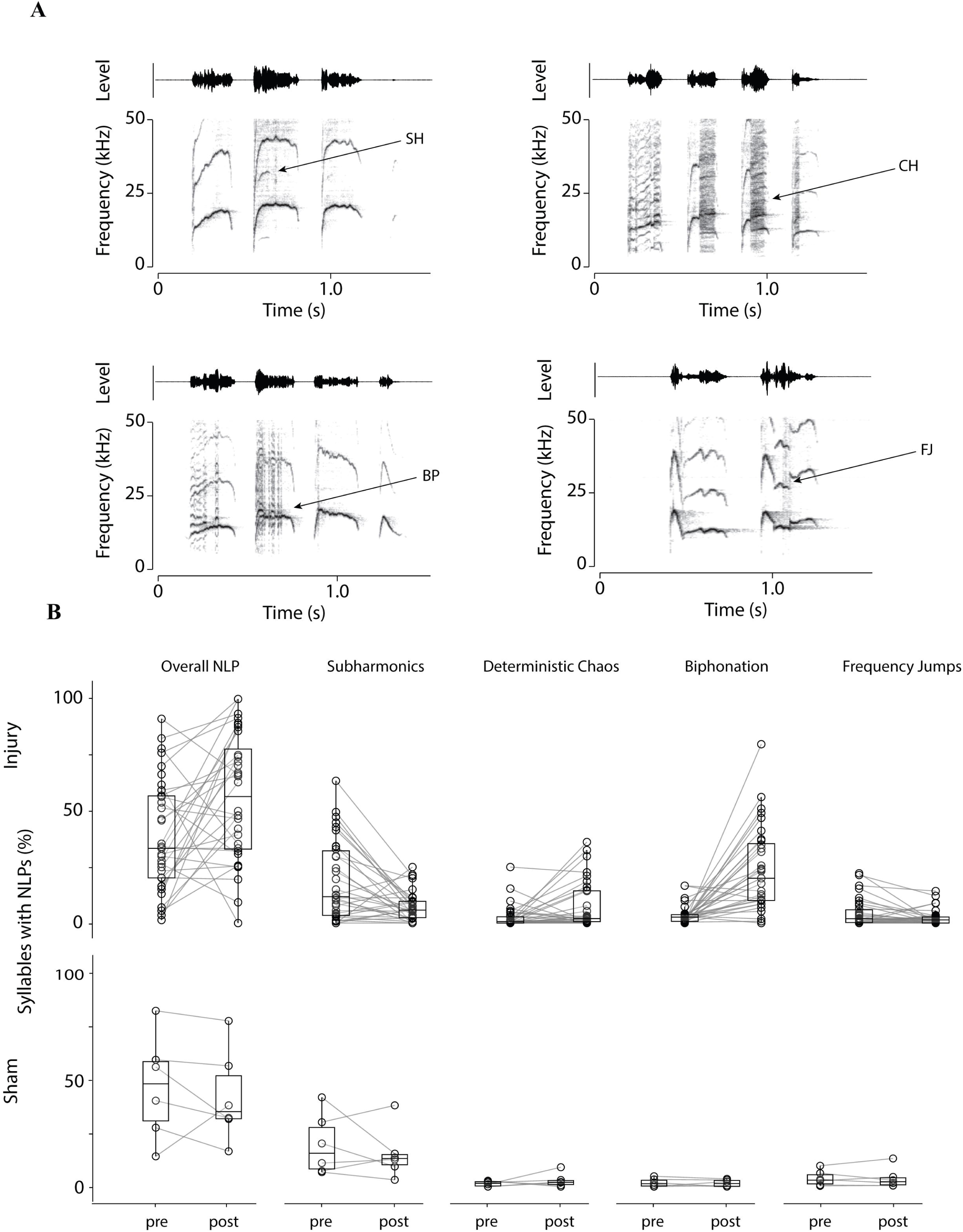
Examples and prevalence of syllables containing nonlinear phenomena (NLPs). (**A**) Spectrograms from four songs illustrating examples of subharmonics (SH), deterministic chaos (CH), biphonation (BP), and frequency jumps (FJ). (**B**) Overall proportion of syllables containing NLPs (%) and the relative occurrence of subharmonics, biphonation, deterministic chaos, and frequency jumps among syllables containing NLPs.

### Characterization of vocal recovery and its association with histological remodeling and vocal load

A common PCA space was established from the mean pre-injury acoustic phenotype of each injured mouse using the seven acoustic variables measured before injury. PCA axes were determined exclusively from these baseline phenotypes; postoperative recordings did not influence the PCA. All postoperative recordings were then projected into this fixed, baseline-derived PCA space. Acoustic recovery was quantified as the Euclidean distance between each postoperative acoustic phenotype and that same mouse’s pre-injury baseline phenotype, such that smaller distances indicated greater recovery toward baseline.

Injured mice showed evidence of vocal recovery over the postoperative period. Across 32 injured mice and 380 mouse-day observations, acoustic distance from each mouse’s pre-injury phenotype decreased significantly with postoperative time in a linear mixed-effects model (β = −0.048 distance units/day, 95% CI −0.081 to −0.014, *p* = 0.018). Thus, on average, mice progressively moved toward their pre-injury acoustic phenotype at a rate of approximately 0.048 distance units per postoperative day. The model included mouse-specific intercepts and recovery slopes, indicating substantial individual variation in recovery trajectories.

Histological severity did not significantly modify the rate of acoustic recovery. The postoperative day × histological score interaction was not significant (β = −0.021 distance units/day per score unit, 95% CI −0.061 to 0.020, *p* = 0.334), providing no evidence that mice with greater histological injury recovered acoustically at a different rate than mice with lower injury scores. Although the negative point estimate was consistent with numerically faster recovery at higher histological scores, the confidence interval included both faster and slower recovery.

Postoperative vocal activity varied substantially among injured mice. Mean vocal activity ranged from 0.07 to 154 calls/day, while total recorded calls ranged from 2 to 3,397 calls per mouse. The duration of postoperative observation also varied, ranging from 6 to 49 calendar days. Mean vocal activity was calculated across all calendar days within each mouse’s observation period, with days without detected calls contributing zero to the mean.

Vocal activity was not associated with overall acoustic distance from baseline. After accounting for postoperative time and histological score, mean postoperative vocal activity was not significantly associated with acoustic distance from baseline (β = 0.039 distance units per 1-SD increase in log-transformed vocal activity, 95% CI −0.494 to 0.572, *p* = 0.886). Thus, mice with greater postoperative vocal activity did not show systematically greater or smaller acoustic deviation from their pre-injury phenotype.

Vocal activity also did not predict the rate of acoustic recovery. The postoperative day × vocal activity interaction was essentially zero (β = −0.00085 distance units/day per 1-SD increase in log-transformed vocal activity, 95% CI −0.033 to 0.031, *p* = 0.960), providing no evidence that greater vocal activity was associated with either faster or slower acoustic recovery.

Overall, the data support recovery following vocal-fold injury, but neither histological severity nor postoperative vocal activity significantly explained differences in the recovery trajectory.

**Figure 5.**
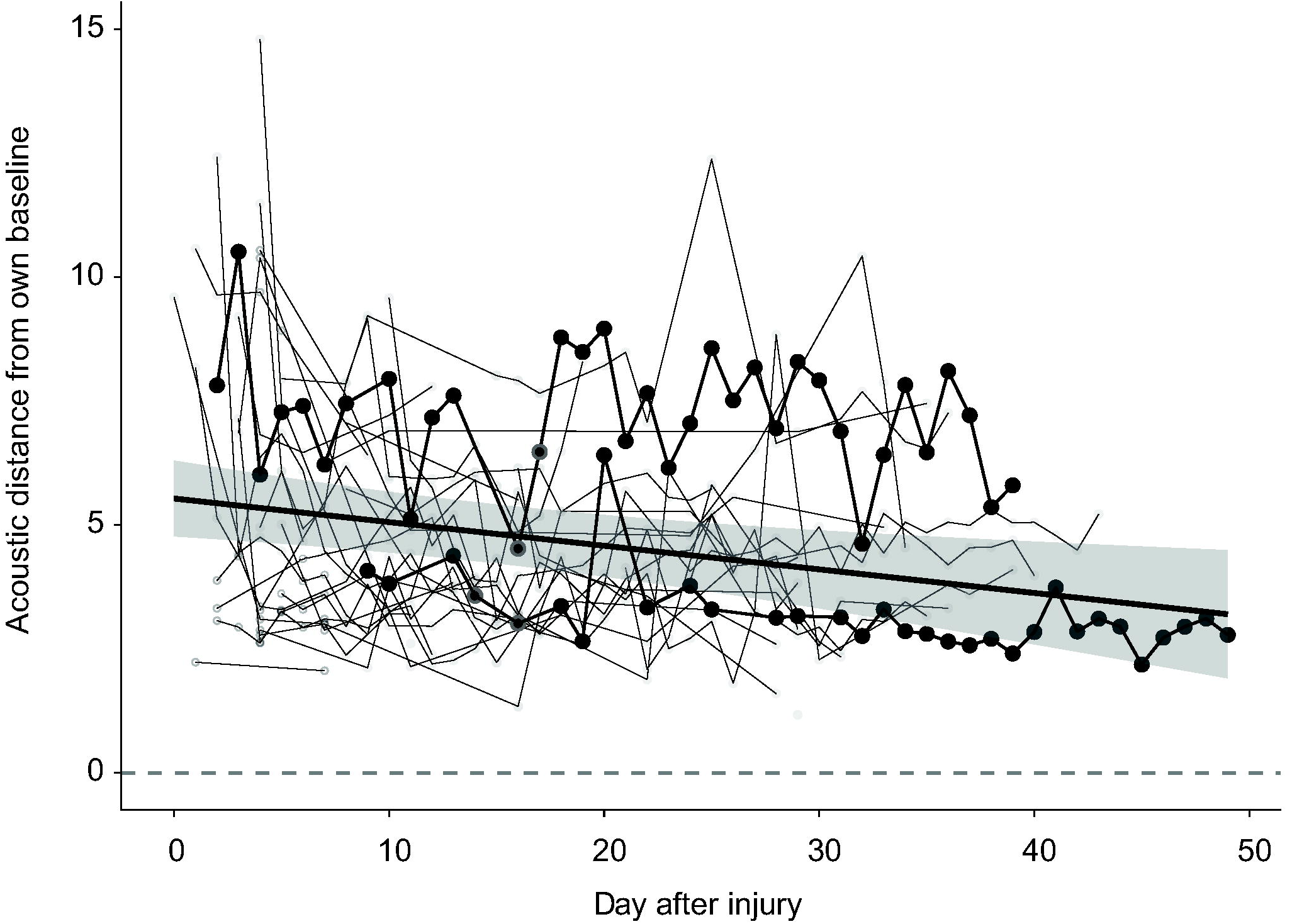
Acoustic recovery following vocal-fold injury. Acoustic distance from each mouse’s own pre-injury acoustic phenotype was calculated in a PCA space defined exclusively by baseline acoustic characteristics. Each gray trajectory represents an individual mouse, with greater distance indicating greater deviation from its pre-injury phenotype. The blue line represents the population-average recovery trajectory estimated from a linear mixed-effects model, with the shaded region indicating the 95% confidence interval. The overall downward trajectory indicates progressive movement toward the pre-injury acoustic phenotype and is consistent with vocal recovery. Two individual mice are highlighted in black as examples of individual recovery trajectories. The horizontal dashed line at zero represents the baseline phenotype.

## Discussion

Predicting the acoustic consequences of structural alterations to the vocal organ remains challenging because vocal fold biomechanics exhibits substantial intrinsic variability (13). Linking vocal phenotypes to specific lesions is difficult because detailed characterization of vocal fold pathology often requires invasive assessment. Our California mouse model of unilateral vocal fold injury illustrates this complexity. Injury altered vocal production, resulting in lower fundamental frequency, reduced vocal intensity, and increased prevalence of nonlinear phenomena. However, neither injury severity at the end of recovery nor vocal activity during recovery was significantly associated with these acoustic changes. Thus, modest structural disruption can alter vocal function without a simple relationship between injury severity and vocal phenotype. Notably, vocal activity remained stable despite acoustic changes, suggesting that injury affected vocal production properties without substantially altering the propensity to vocalize. Below, we consider how stable vocal activity alongside altered acoustic structure informs relationships between vocal behavior and the mechanical properties of the vocal apparatus.

### Stable vocal activity

Average daily vocal activity was unchanged following unilateral vocal fold injury. Although injury produced localized tissue damage, inflammation, and subsequent structural remodeling, it did not reduce spontaneous vocal activity, suggesting that localized injury was insufficient to produce generalized suppression of vocal behavior.

Local tissue injury should be distinguished from sickness behavior, a coordinated set of physiological and behavioral adaptations that occur during systemic immune activation and is mediated by interactions between peripheral immune signals and the central nervous system (14–16). The maintenance of vocal activity suggests that localized injury did not elicit a sufficiently strong systemic or behavioral response to suppress vocalization.

Two injured animals produced no vocalizations after injury. One exhibited a large injury extending deep into the vocal fold (Supplementary Fig. 1c), whereas the other showed no apparent vocal-fold abnormalities after four weeks of healing. These cases suggest that vocal activity can be affected in some animals, but absence of vocalization is not necessarily explained by apparent pathology. Overall, preserved vocal activity despite altered acoustic features enables noninvasive investigation of injury effects on vocal production.

### Altered acoustic parameters and nonlinear phenomena

Fundamental frequency depends on vocal fold biomechanics, subglottal pressure, vocal fold length and tension, and neuromuscular control (17). Its reduction following injury therefore likely reflects altered biomechanics: damage to muscle and lamina propria can change tissue mechanics and vibration, while unilateral damage may disrupt vibratory symmetry. Similar effects occur in humans and vocal-fold models (19–21).

Reduced vocal intensity suggests decreased sound-production efficiency, potentially from incomplete glottal closure and increased air leakage, which reduce aerodynamic-to-vibratory energy transfer (22–26), consistent with weaker output in unilateral human vocal-fold pathology (27–28). However, reduced motivation to vocalize or lower subglottal pressure during vocalization must also be considered as alternative explanations.

Injured mice also produced shorter syllables at higher repetition rates while maintaining song structure (syllables/song), indicating altered temporal organization. Shorter syllables may reflect unstable oscillation from increased asymmetry/reduced efficiency or altered sensory feedback; faster repetition may compensate for reduced syllable duration or result from premature syllable termination and earlier onset of subsequent vocalizations. Similar temporal adaptations occur in human dysphonia (29), suggesting injury alters vocal motor control beyond spectral and intensity changes.

Nonlinear phenomena provide further evidence for airflow-driven vocal fold vibration rather than aerodynamic whistling. Although more syllables contained nonlinear phenomena, subharmonics decreased while biphonation and deterministic chaos increased; frequency jumps were unchanged. Increased biphonation and chaos indicate greater vibratory instability, as observed in pathological human voices (30–31). However, the inverse relationship among subharmonics, biphonation, and deterministic chaos has not previously been reported.

### Biological significance

The vocal fold injury observed here involved rupture of a vocal fold blood vessel and blood extravasation extending primarily through the lamina propria and thyroarytenoid (TA) muscle, with preserved epithelium. This pattern is consistent with acute vocal fold hemorrhage caused by submucosal vessel rupture and blood accumulation (32). Because the lamina propria and TA muscle contribute to the mechanical properties and vibration of the vocal folds, injury and subsequent remodeling of these tissues provide a plausible substrate for the persistent changes in vocal acoustics observed here.

The most striking finding was that vocal activity remained stable despite substantial changes in acoustic parameters. Thus, continued performance of the behavior did not indicate preserved function of the tissues underlying sound production. Instead, animals continued to vocalize while producing sounds with lower fundamental frequency and intensity, shorter syllables, and more frequent nonlinear phenomena. These findings demonstrate that behavioral output can remain intact while the physiological and biomechanical machinery supporting that behavior is substantially altered (33). Natural vocalizations may therefore provide a sensitive readout of otherwise cryptic changes in tissue function that would not be apparent from vocal activity alone.

The increased occurrence of nonlinear phenomena further indicates that injury altered the dynamics of vocal-fold vibration rather than simply reducing vocal output. Such phenomena can arise from vocal-fold instabilities, suggesting that acoustic variation may reflect changes in biomechanical state rather than solely active vocal modulation.

If vocal-fold injury occurs naturally through trauma, repeated vocal loading, or disease, it could contribute to transient variation in vocal phenotype by altering acoustic output without suppressing vocal activity. More broadly, our findings illustrate how tissue injury and remodeling can reshape the phenotype of a behavior even when the behavior itself is maintained.

### Translational significance

Our findings raise questions about how continued vocal use interacts with tissue recovery. Voice rest is an important component of early vocal fold injury management and is thought to promote wound healing (34, 35), yet the relationship between vocal loading and recovery remains incompletely understood. California mice continued vocalizing after injury, and variation in vocal activity did not explain persistence of acoustic alterations. Thus, within the range of vocal use observed here, vocal activity alone was not a reliable indicator of post-injury vocal recovery. Experimental manipulation of vocal loading will be necessary to determine whether continued use influences tissue remodeling or prolongs functional impairment.

The California mouse provides a useful model for addressing these questions because, like humans, it produces vocalizations through airflow-induced, self-sustained vocal fold vibration followed by dynamic vocal-tract filtering (7, 8). Natural vocalizations can also be recorded repeatedly and noninvasively from the same individuals, allowing vocal phenotype to be tracked alongside tissue changes over time. This framework could be extended to other injury paradigms and interventions to test how vocal use and tissue remodeling interact during recovery.

## Conclusions

Impairment to the vocal folds can substantially alter acoustic phenotype while leaving vocal activity largely intact, demonstrating that persistence of a behavior does not necessarily indicate preservation of the biomechanical systems underlying its performance. Natural vocalizations can therefore provide a sensitive, noninvasive readout of otherwise cryptic changes in the physical substrate of behavior. More broadly, this framework provides a tractable approach for linking microscopic tissue injury and remodeling to longitudinal changes in behavioral phenotype and for testing how mechanical loading shapes behavioral recovery.

