## Supplemental Figures for "Vocal fold injury alters acoustic phenotype despite preserved vocal activity"

Supplemental Material

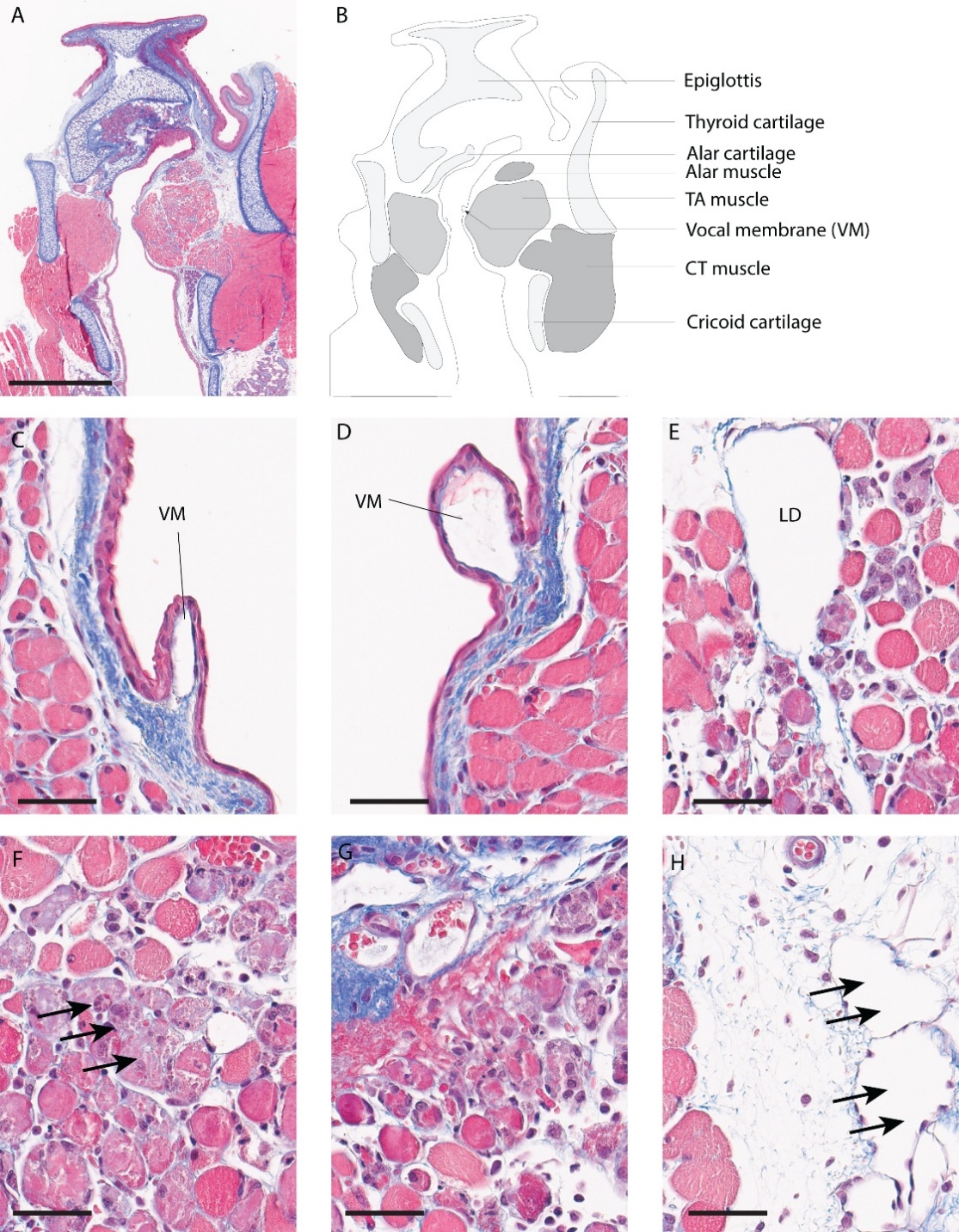

**Supplemental Figure 1a: Representative histological section through the mid-membranous vocal folds of a male California mouse with a histopathology score of 5 (maximum possible score = 9; highest score in this study: 6).** Sections were stained with Masson's trichrome. (**A**, **B**) Low-magnification overview of the larynx. Higher-magnification images of the left (C) and right (**D**) membranous vocal folds. Note the dilated lymphatic vessel within the injured (right) vocal fold. VM, vocal membrane. (**E**) Lymphatic dilation (LD) within multiple degenerated myofibers of the thyroarytenoid (TA) muscle. (**F**) Multiple degenerated myofibers (arrows) with few scattered lymphocytes. (**G**) Multiple degenerated, necrotic, and atrophic myofibers with few scattered lymphocytes. (**H**) Lymphatic dilation within the connective tissue between the lateral TA muscle and the thyroid cartilage. Scale bars = 100 μm in all panels.

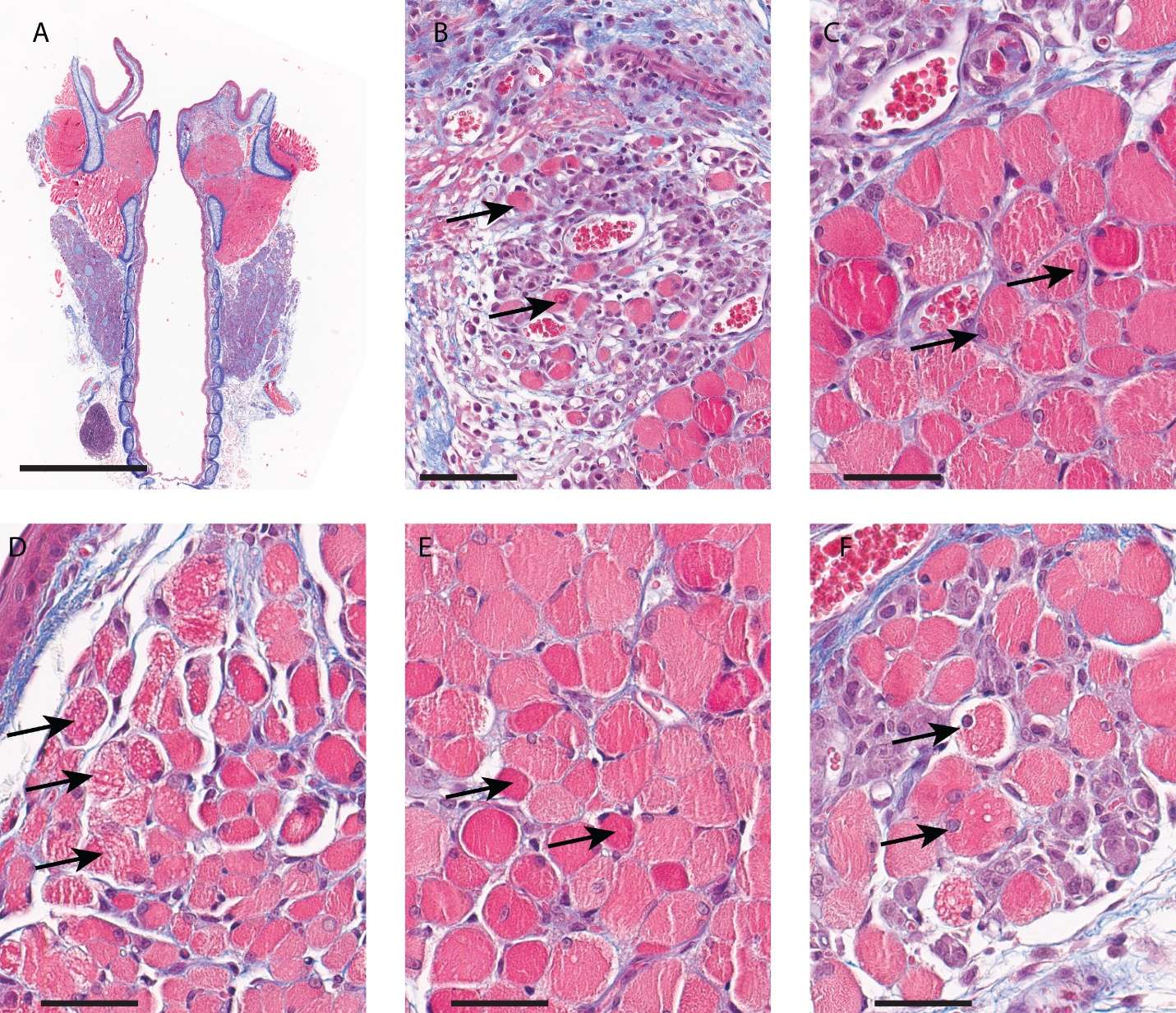

**Supplemental Figure 1b: Representative histological section through the mid-membranous vocal folds of a male California mouse with a histopathology score of 6 (maximum possible score = 9; highest score observed in this study = 6).** Sections were stained with Masson’s trichrome. (**A**) Low-magnification overview of the larynx. (**B**) Higher-magnification image shows large areas of degenerated, necrotic and atrophic myofibers characterized by atrophic and hypereosinophilic myofibers (arrow). (**C**) Swollen degenerating myofibers have reactive/ plump nuclei (arrow). (**D**) Multiple degenerating and atrophic myofibers contain multiple cytoplasmic vacuoles (arrow). (**E**) Multiple swollen and degenerating myofibers show hypereosinophilic cytoplasm. (**F**) Multiple swollen and degenerating myofibers contain several cytoplasmic vacuoles and reactive/plump nuclei. Scale bars = 100 μm in all panels.

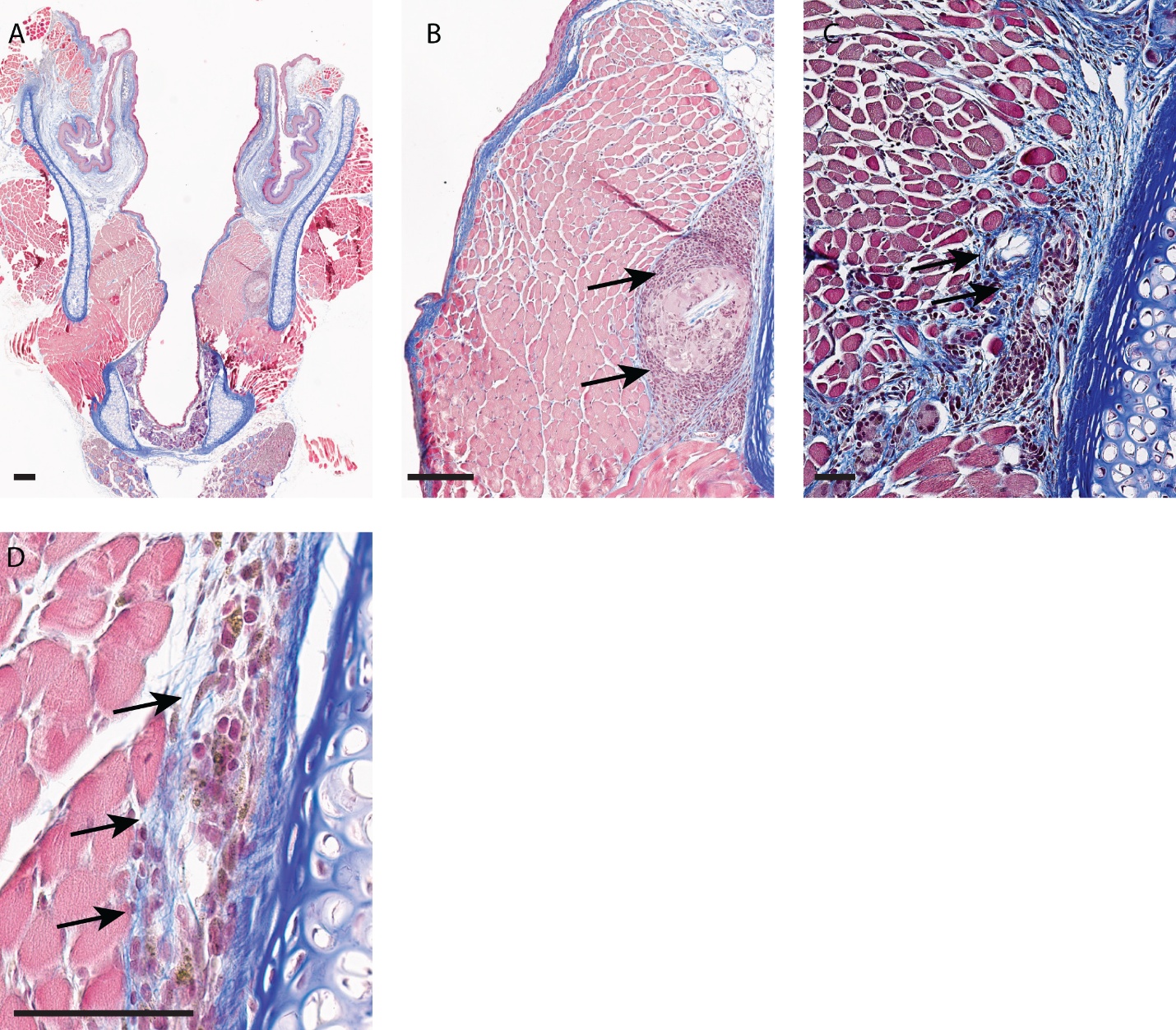

**Supplemental Figure 1c: Histological section through the mid-membranous vocal folds of a male California mouse with a histopathology score of 4 after 4 weeks of recovery**. This animal was one of two animals that did not vocalize following vocal fold injury. Sections were stained with Masson's trichrome. (**A**) Low-magnification overview of the larynx. (**B**–**D**) Necrotic tissue embedded within the thyroarytenoid muscle near its boundary with the thyroid cartilage. Arrows indicate areas of necrosis and increased fibrosis. Scale bars = 100 μm in all panels.

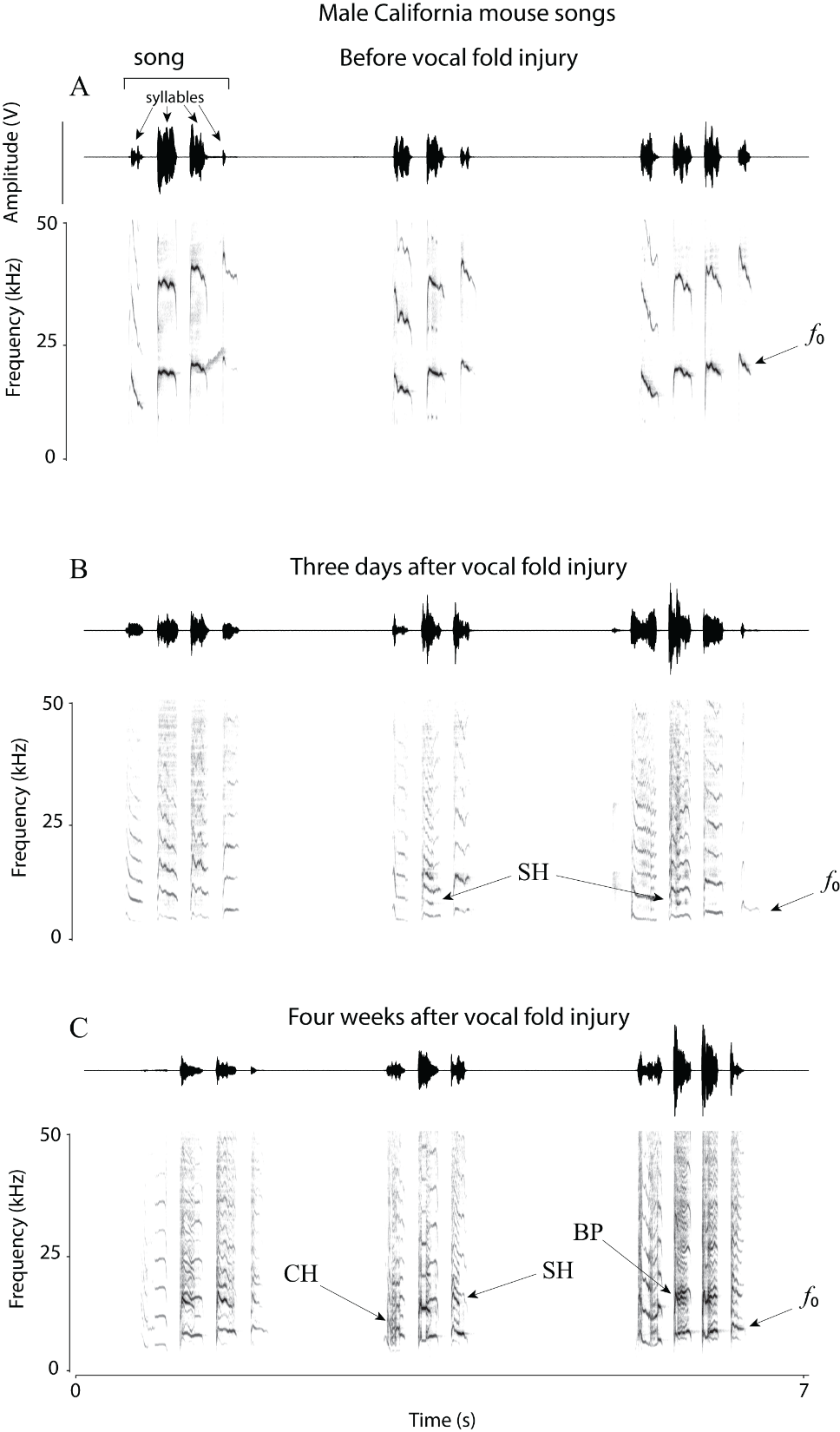

**Supplemental Figure 2a**: **Representative songs of a California mouse before and after unilateral vocal fold injury.** Each panel shows three songs from the same male recorded before injury (**A**) and after injury (**B**, **C**). Vocalizations are displayed as waveforms (top) and spectrograms (bottom). Examples of nonlinear phenomena are indicated, including deterministic chaos (CH), biphonation (BP), and subharmonics (SH). *f*_0_, fundamental frequency.

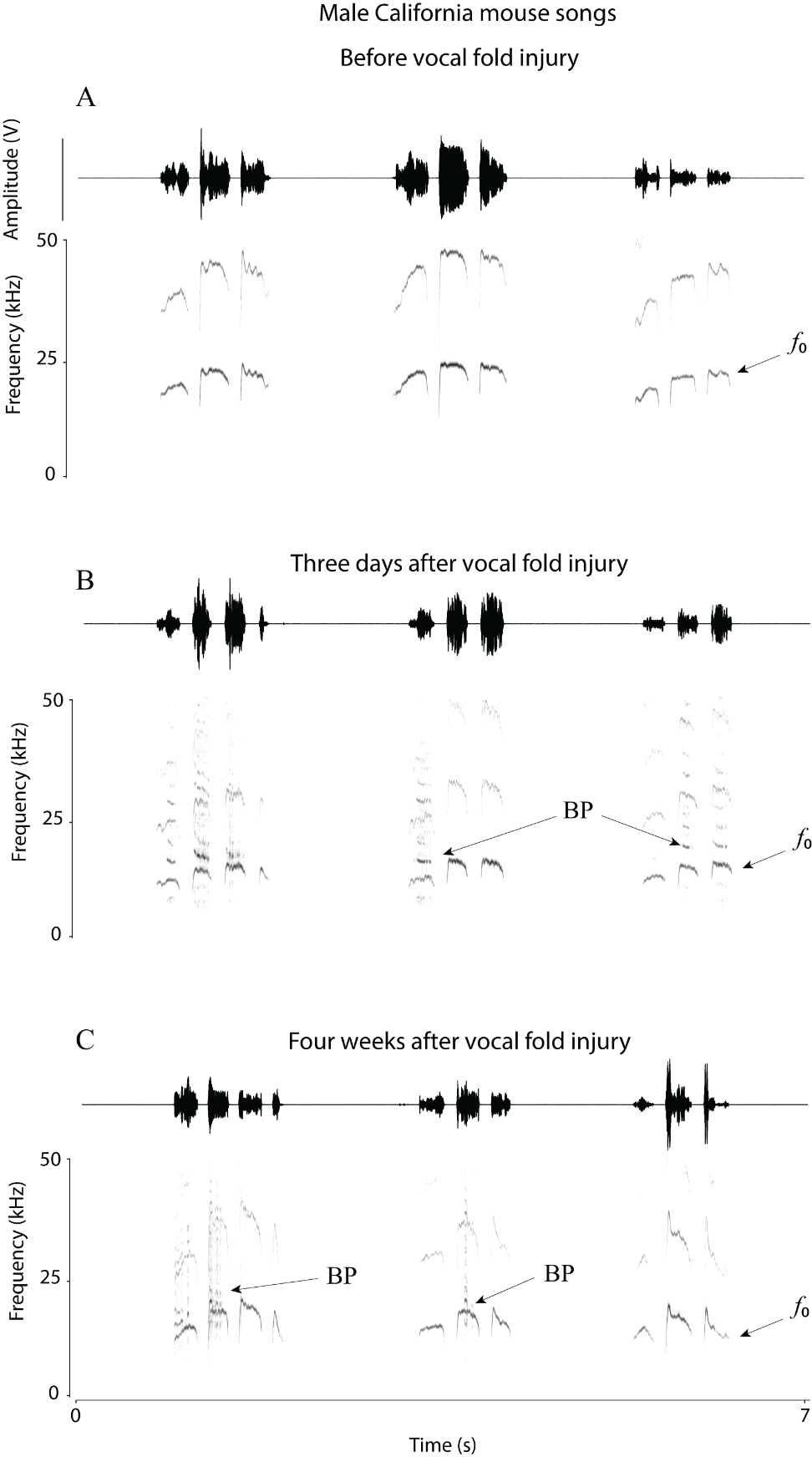

**Supplemental Figure 2b**: Songs of California mice before and after vocal fold injury. Each panel shows three songs from the same male mouse before (**A**) and after (**B**, **C**) injury. Sound is depicted as a waveform and spectrogram. This animal exhibited mostly biphonation (BP) after injury. *f*_0_, fundamental frequency.

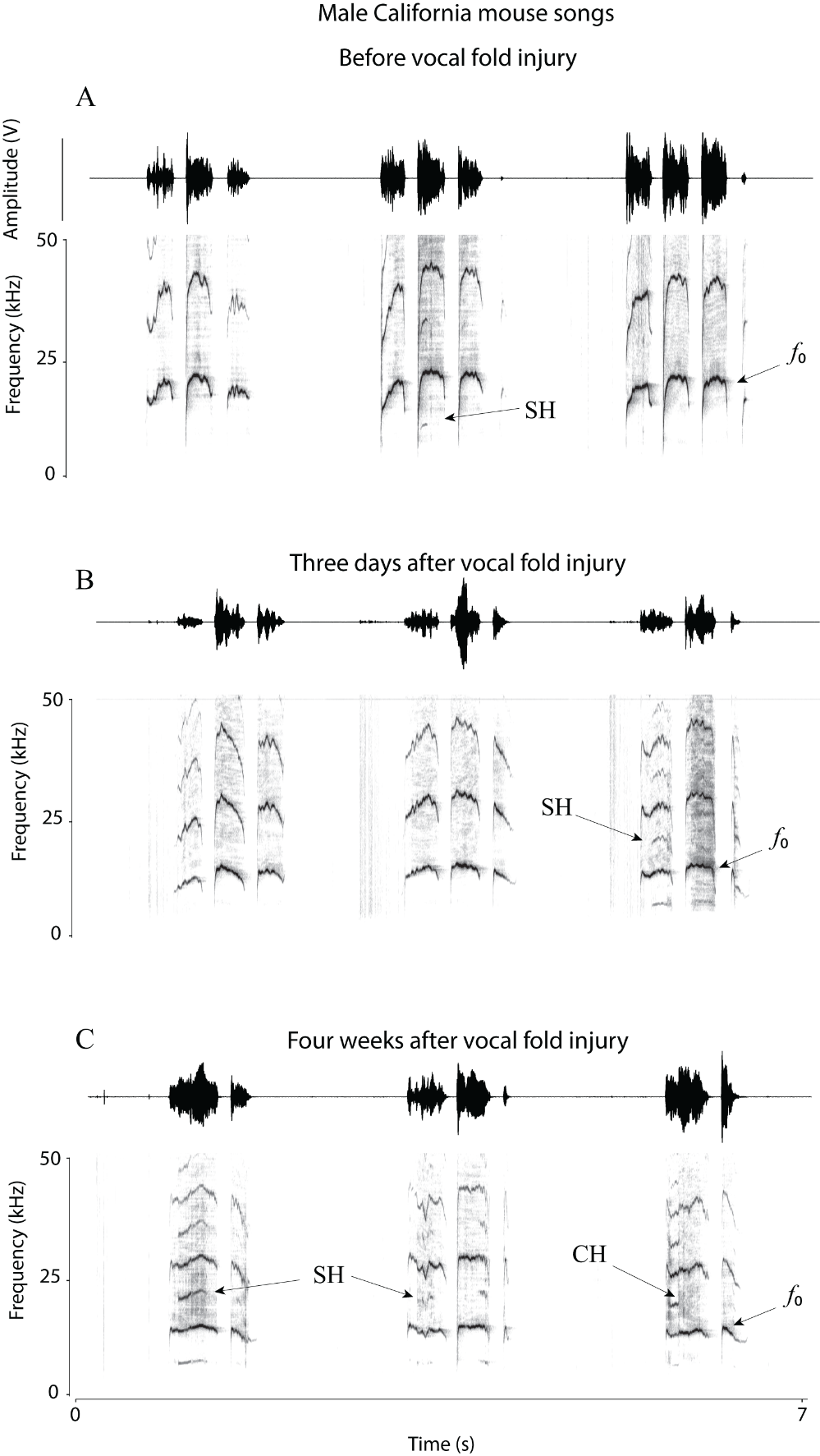

**Supplemental Figure 2c**: Songs of California mice before and after vocal fold injury. Each panel shows three songs from the same male mouse before (**A**) and after (**B**, **C**) injury. Sound is depicted as a waveform and spectrogram. This animal exhibited deterministic chaos (CH) and subharmonics (SH) after injury. Furthermore mean syllable fundamental frequency (*f*_0Mean_) decreased from about 20 kHz to 14 kHz after injury. *f*_0_, fundamental frequency.
